# Comprehensive analysis of RNA-sequencing to find the source of 1 trillion reads across diverse adult human tissues

**DOI:** 10.1101/053041

**Authors:** Serghei Mangul, Harry Taegyun Yang, Nicolas Strauli, Franziska Gruhl, Hagit T. Porath, Kevin Hsieh, Linus Chen, Timothy Daley, Stephanie Christenson, Agata Wesolowska-Andersen, Roberto Spreafico, Cydney Rios, Celeste Eng, Andrew D. Smith, Ryan D. Hernandez, Roel A. Ophoff, Jose Rodriguez Santana, Erez Y. Levanon, Prescott G. Woodruff, Esteban Burchard, Max A. Seibold, Sagiv Shifman, Eleazar Eskin, Noah Zaitlen

**Affiliations:** Department of Computer Science, University of California Los Angeles, Los Angeles, USA; Institute for Quantitative and Computational Biosciences, University of California Los Angeles, Los Angeles, USA; Biomedical Sciences Graduate Program, University of California, San Francisco, CA, USA; Center for Integrative Genomics, University of Lausanne, Lausanne, Switzerland; Swiss Institute of Bioinformatics, Lausanne, Switzerland; The Mina and Everard Goodman Faculty of Life Sciences, Bar-Ilan University, Ramat-Gan, Israel; Department of Bioengineering, University of California Los Angeles, Los Angeles, USA; Department of Molecular and Computational Biology, University of Southern California, CA, USA; Division of Pulmonary, Critical Care, Sleep and Allergy, Department of Medicine, and Cardiovascular Research Institute, University of California, San Francisco, CA, USA; Center for Genes, Environment, and Health, National Jewish Health, Denver, CO, USA; Department of Medicine, University of California, San Francisco, CA, USA; Department of Bioengineering and Therapeutic Sciences, University of California, San Francisco, CA, USA; Institute for Quantitative Biosciences, University of California, San Francisco, CA, USA; Institute for Human Genetics, University of California San Francisco, San Francisco, CA, USA; Center for Neurobehavioral Genetics, Semel Institute for Neuroscience and Human Behavior, University California Los Angeles, Los Angeles, USA; Department of Human Genetics, University of California Los Angeles, Los Angeles, USA; Department of Psychiatry, Brain Center Rudolf Magnus, University Medical Center Utrecht, Utrecht, The Netherlands; Pediatric Pulmonology, San Juan, Puerto Rico; Department of Pediatrics, National Jewish Health, Denver, CO, USA; University of Colorado School of Medicine, Denver CO, USA; Department of Genetics, The Institute of Life Sciences, The Hebrew University of Jerusalem, Jerusalem, Israel

## Abstract

High throughput RNA sequencing technologies have provided invaluable research opportunities across distinct scientific domains by producing quantitative readouts of the transcriptional activity of both entire cellular populations and single cells. The majority of RNA-Seq analyses begin by mapping each experimentally produced sequence (i.e., read) to a set of annotated reference sequences for the organism of interest. For both biological and technical reasons, a significant fraction of reads remains unmapped. In this work, we develop Read Origin Protocol (ROP) to discover the source of all reads originating from complex RNA molecules, recombinant T and B cell receptors, and microbial communities. We applied ROP to 8,641 samples across 630 individuals from 54 tissues. A fraction of RNA-Seq data (n=86) was obtained in-house; the remaining data was obtained from the Genotype-Tissue Expression (GTEx v6) project. To generalize the reported number of accounted reads, we also performed ROP analysis on thousands of different, randomly selected, and publicly available RNA-Seq samples in the Sequence Read Archive (SRA). Our approach can account for 99.9% of 1 trillion reads of various read length across the merged dataset (n=10641). Using in-house RNA-Seq data, we show that immune profiles of asthmatic individuals are significantly different from the profiles of control individuals, with decreased average per sample T and B cell receptor diversity. We also show that immune diversity is inversely correlated with microbial load. Our results demonstrate the potential of ROP to exploit unmapped reads in order to better understand the functional mechanisms underlying connections between the immune system, microbiome, human gene expression, and disease etiology. ROP is freely available at <u>https://github.com/smangul1/rop</u> and currently supports human and mouse RNA-Seq reads.

## INTRODUCTION

Advances in RNA sequencing (RNA-seq) technology have provided an unprecedented opportunity to explore gene expression across individual, tissues, and environments (Cloonan et al., 2008; Sultan et al., 2008; Tang et al., 2009) by efficiently profiling the RNA sequences present in a sample of interest (Z. Wang, Gerstein, & Snyder, 2009). RNA-seq experiments currently produce tens of millions of short read subsequences sampled from the complete set of RNA transcripts that are provided to the sequencing platform. An increasing number of bioinformatic protocols are being developed to analyze reads in order to annotate and quantify the sample’s transcriptome (Mihaela Pertea, 2015; Nicolae, Mangul, Mandoiu, & Zelikovsky, 2011; Trapnell et al., 2010). When a reference genome sequence or, preferably, a transcriptome of the sample is available, mapping-based RNA-seq analysis protocols align the RNA-seq reads to the reference sequences, identify novel transcripts, and quantify the abundance of expressed transcripts.

Unmapped reads, the reads falling to map to the human reference, are a large and often overlooked output of standard RNA-seq analyses. Even in carefully executed experiments, the *unmapped reads* can comprise a substantial fraction of the complete set of reads produced; for example, approximately 9%-20% of reads are unmapped in recent large human RNA-seq projects (Ardlie et al., 2015; Li, Tighe, Nicolet, Grove, Levy, Farmerie, Viale, Wright, Schweitzer, Gao, Kim, et al., 2014; Seqc/Maqc-Iii Consortium, 2014). Unmapped reads can arise due to technical sequencing artifacts that were produced by low quality and error prone copies of the nascent RNA sequence being sampled (Ozsolak & Milos, 2011). A recent study by Baruzzo et al., (2017) suggests that at least 10% of the reads simulated from human references remain unmapped across 14 contemporary state-of-the art RNA alligners. This rate may be due to shortcomings of the aligner’s efficient yet heuristic algorithms (Siragusa, Weese, & Reinert, 2013). Reads can also remain unmapped due to unknown transcripts (Grabherr et al., 2011), recombined B and T cell receptor sequences (Blachly et al., 2015; N. B. Strauli & Hernandez, 2016), A-to-G mismatches from A-to-I RNA editing (Porath, Carmi, & Levanon, 2014), trans-splicing (Wu et al., 2014), gene fusion (X.-S. Wang et al., 2009), circular RNAs (Jeck & Sharpless, 2014), and the presence of non-host RNA sequences (Kostic et al., 2011) (e.g., bacterial, fungal, and viral organisms).

In this work, we report the development of a comprehensive method that can characterize the origin of unmapped reads obtained by RNA-seq experiments. Analyzing unmapped reads can inform future development of read mapping methods, provide access to additional biological information, and resolve the irksome puzzle of the origin of unmapped reads. We developed the Read Origin Protocol (ROP), a multi-step approach that leverages accurate alignment methods for both host and microbial sequences. The ROP tool contains a combination of novel algorithms and existing tools focused on specific categories of *unmapped reads* (Blachly et al., 2015; Brown, Raeburn, & Holt, 2015; Chuang et al., 2015; Kostic et al., 2011; N. Strauli & Hernandez, 2015). The comprehensive analytic nature of the ROP tool prevents biases that can otherwise arise when using standard targeted analyses. Currently, ROP supports human and mouse RNA-Seq data.

## RESULTS

### ROP – a computational protocol to explain unmapped reads in RNA-Sequencing

Mapping-based RNA-seq analysis protocols overlook reads that fail to map onto the human reference sequences (i.e., *unmapped reads*). We designed a read origin protocol (ROP) that identifies the origin of both mapped and unmapped reads (Fig. 1). The protocol first identifies human reads by mapping them onto a reference genome and transcriptome using a standard high-throughput mapping algorithm (Kim et al., 2013). We used tophat v. 2.0.12 with ENSEMBL GRCh37 transcriptome and hg19 build, but many other mapping tools are available and have recently been reviewed by Baruzzo et al., 2017). After alignment, reads are grouped into genomic (e.g., CDS, UTRs, introns) and repetitive (e.g., SINEs, LINEs, LTRs) categories. The rest of the ROP protocol characterizes the remaining *unmapped reads*, which failed to map to the human reference sequences.

**Figure 1.**
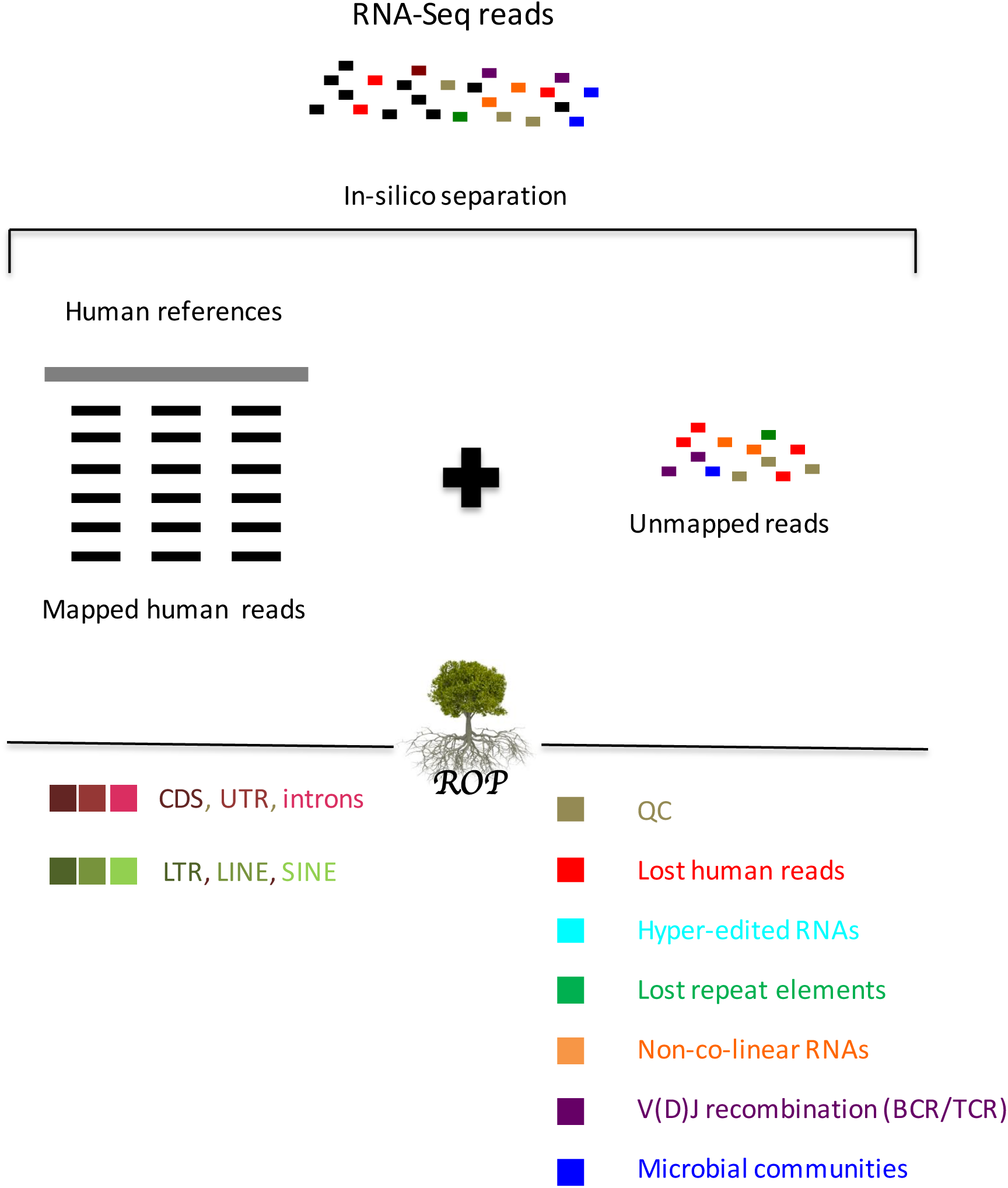
Schematic of the Read Origin Protocol (ROP). Human reads are identified by mapping all reads onto the reference sequences using a standard high-throughput mapping algorithm. ROP protocol categorizes mapped reads into genomic (red colors) and repetitive (green colors) categories. Unmapped reads that fail to map are extracted and further filtered to exclude low quality reads, low complexity reads, and reads from rRNA repeats (brown color). ROP protocol is able to identify unmapped reads aligned to human references with use of a more sensitive alignment tool (lost human reads: red color), unmapped reads aligned to human references with excessive (‘hyper’) editing (hyper-edited RNAs: cyan color), unmapped reads aligned to the repeat sequences (lost repeat elements: green color), unmapped reads spanning sequences from distant loci (non-co-linear: orange color), unmapped reads spanning antigen receptor gene rearrangement in the variable domain (V(D)J recombination of BCR and TCR: violet color), and unmapped reads aligned to the microbial reference genomes and marker genes (microbial reads: blue color).

The ROP protocol effectively processes the *unmapped reads* in seven steps. First, we apply a quality control step to exclude low-quality reads, low-complexity reads, and reads that match rRNA repeat units among the unmapped reads (FASTQC (Andrews & others, 2010), SEQCLEAN (“https://sourceforge.net/projects/seqclean/,” n.d.)). Next, we employ Megablast (Camacho et al., 2009), a more sensitive alignment method, to search for human reads missed due to heuristics implemented for computational speed in conventional aligners and reads with additional mismatches. These reads typically include those with mismatches and short gaps relative to the reference set, but they can also include perfectly matched reads. Hyper-editing pipelines recognize reads with excessive (‘hyper’) editing, which are usually rejected by standard alignment methods due to many A-to-G mismatches (Porath, Carmi, & Levanon, 2014). We use a database of repeat sequences to identify lost repeat reads among the *unmapped reads*. Megablast, and similar sensitive alignment methods, are not designed to identify ‘non-co-linear’ RNA (Chuang et al., 2015) reads from circRNAs, gene fusions, and trans-splicing events, which combine a sequence from distant elements. For this task, we independently map 20bp read anchors onto the genome (see Supplemental Methods). Similarly, reads from BCR and TCR loci, which are subject to recombination and somatic hyper-mutation (SHM), require specifically designed methods. For this case, we use IgBlast (Ye, Ma, Madden, & Ostell, 2013). The remaining reads that did not map to any known human sequence are potentially microbial in origin. We use microbial genomes and phylogenetic marker genes to identify microbial reads and assign them to corresponding taxa (Truong et al., 2015). Microbial reads can be introduced by contamination or natural microbiome content in the sample, such as viral, bacterial, fungi, or other microbial species (Salter et al., 2014).

Taken together, ROP considers six classes of *unmapped reads*: (1) lost human reads, (2) hyper-edited reads, (3) lost repeat elements, (4) reads from ‘non-co-linear’ (NCL) RNAs, (5) reads from the recombination of BCR and TCR segments (i.e. V(D)J recombination), and (6) microbial reads. Previously proposed individual methods do examine some of these classes (Blachly et al., 2015; Brown et al., 2015; Chuang et al., 2015; Kostic et al., 2011; N. Strauli & Hernandez, 2015). However, we find that performing a sequential analysis, in the order described above, is critical for minimizing misclassification of reads due to homologous sequences between the different classes. Furthermore, as shown in the Results section below, only a comprehensive analysis allows comparison across these classes. We have demonstrated the robustness of the proposed approach against alternating order of steps and values of the thresholds (Supplemental Methods and Supplemental Methods Figure SM1). Complete details of ROP, including all parameters and thresholds used, are provided in the Supplemental Methods.

### The ROP protocol is able to account for 99.9% of all reads

To test ROP, we applied it to one trillion RNA-Seq reads across 54 tissues from 2630 individuals. The data was combined from 3 studies: (1) in-house RNA-Seq data (n=86) from the peripheral blood, nasal, and large airway epithelium of asthmatic and control individuals (S1); (2) multi-tissue RNA-Seq data from Genotype-Tissue Expression (GTEx v6) from 53 human body sites (Consortium & others, 2015) (n=8555) (S2); (3) randomly selected RNA-Seq samples from the Sequence Read Archive (SRA) (n=2000) (S3). Unless otherwise noted, we reported percentage of reads averaged across 3 datasets.

RNA-Seq data obtained from the three sources represent a large collection of tissue types and read diversity. We selected these three sources to most accurately model the precision and broad applicability of ROP. The in-house RNA-Seq data was collected from 53 asthmatics and 33 controls. RNA-seq libraries were prepared from total RNA with two types of RNA enrichment methods: (1) Poly(A) enrichment libraries, applied to RNA from peripheral blood and nasal epithelium (n=38), and (2) ribo-depletion libraries, applied to RNA from large airway epithelium (n=49). The GTEx dataset was derived from 38 solid organ tissues, 11 brain subregions, whole blood, and three cell lines across 544 individuals. Randomly selected SRA RNA-Seq samples included samples from whole blood, brain, various cell lines, muscle, and placenta. Length of reads from in-house data was 100bp, read length in Gtex data was 76bp, read length in SRA data ranged from 36bp to 100bp. In total, 1 trillion reads (97 Tbp) derived from 10641 samples were available for ROP (Supplemental Table S1 and Supplementary Methods). For counting purposes, the pairing information of the reads is disregarded, and each read from a pair is counted separately.

We used standard read mapping procedures to obtain mapped and unmapped reads from all three data sources. Read mapping for GTEx data was performed by the GTEx consortium using TopHat2 (Kim et al., 2013). Following the GTEx consortium practice, we used TopHat2 to map reads from in-house and SRA studies. High-throughput mapping using TopHat2 (Kim et al., 2013) recovered 83.1% of all reads from three studies (Fig. 2.a), with the smallest fraction of reads mapped in the SRA study (79% mapped reads). From the *unmapped reads*, we first excluded low-quality/low-complexity reads and reads mapping to the rRNA repeating unit, which together accounted for 7.0% and 2.4% of all reads, respectively (Fig. 2.b). We were then able to align unmapped reads to human reference sequences (5.7% of all reads, Fig. 2.c) and identify “hyper-edited” reads (0.1% of all reads Fig. 2.d). We then referenced repeat sequences (0.2% of all reads, Fig. 2.d), reads identified as ‘non-co-linear’(NCL) RNAs (circRNAs, gene fusion or trans-splicing) (0.3% of all reads, Fig. 2.e), and reads mapped to recombined B and T cell receptors (0.02% off all reads, Fig. 2.f). The remaining reads were mapped to the microbial sequences (1.4% off all reads, Fig. 2.g). Following the seven steps of ROP, the origins of 99.9% of reads were identified. Genomic profile of unmapped reads for each dataset is separately reported in Table S2. Uncategorized reads from SRA samples are freely available at https://smangul1.github.io/recycle-RNA-seq/. This resource allows the bioinformatics community to further increase the number of reads with known origin.

**Figure 2.**
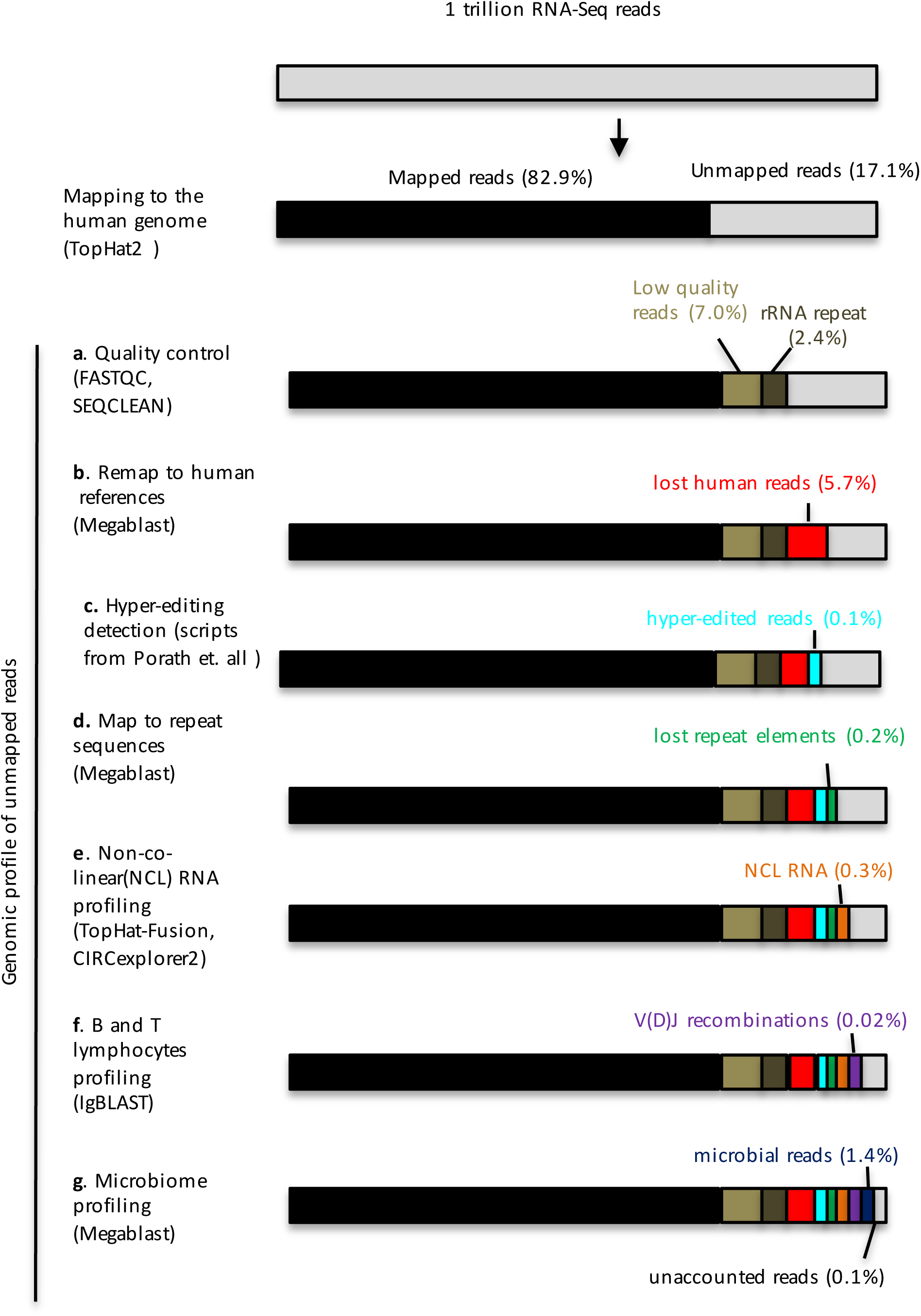
Genomic profile of unmapped reads across 10641 samples and 54 tissues. Percentage of unmapped reads for each category is calculated as a fraction from the total number of reads. Bars of the plot are not scaled. Human reads (black color) mapped to reference genome and transcriptome via TopHat2. (a) Low quality/low-complexity (light brown) and reads matching rRNA repeating unit (dark brown) were excluded. (b) Hyper-edited reads are captured by hyper-editing pipeline proposed in (Porath et al., 2014). (c) ROP identifies lost human reads (red color) from unmapped reads using a more sensitive alignment. (d) ROP identifies lost repeat sequences (green color) by mapping unmapped reads onto the reference repeat sequences. (e) Reads arising from trans-spicing, gene fusion and circRNA events (orange color) are captured by a TopHat-Fusion and CIRCexplorer2 tools. (f) IgBlast is used to identify reads spanning B and T cell receptor gene rearrangement in the variable domain (V(D)J recombinations) (violet color). (g) Microbial reads (blue color) are captured by mapping the reads onto the microbial reference genomes.

### The ROP protocol identifies lost human reads

Some human reads may remain unmapped due to the heuristic nature of high throughput aligners (Baruzzo et al., 2016; Siragusa et al., 2013). As shown by Baruzzo et al., even the best performing RNA-Seq aligners fail to map at least 10% of reads simulated from the human references. To prevent misclassification of reads derived from human genome into other downstream ROP categories, we used the slower and more sensitive Megablast aligner on this subset of unmapped reads. This method allows us to filter an additional 5.7% of human reads.

We investigated the impact of mapping parameters and RNA-Seq aligners on the number of unmapped reads. We additionally used STAR (Dobin et al., 2013) and added results for sensitive and very sensitive mapping settings of each of the tools (Supplemental Methods and Supplemental Table S4). We observe that an alternative aligner and a more sensitive mapping setting has no substantial effect on the number of mapped reads (Supplementary Table S5). This is in line with Baruzzo et al., 2016, which have shown that optimizing the parameters of RNA-Seq aligner is a non-trivial task and methods with good performance for the default setting is a preferred choice.

Using both mapped and unmapped reads across the studies, we classified on average 7.5% of the RNA-Seq reads as repetitive sequences originated from various repeat classes and families (Supplemental Fig. S2). We observe Alu elements to have 33% relative abundance, which was the highest among all the repeat classes. Among DNA repeats, hAT-Charlie was the most abundant element with 50% relative abundance (Supplemental Fig. S3). Among SVA retrotransposons, SVA-D was the most abundant element with 50% relative abundance (Supplemental Fig. S4). Consistent with repEnrich (Criscione, Zhang, Thompson, Sedivy, & Neretti, 2014), when using in-house data we observe the differences in proportions of L1 and Alu elements between poly(A) and ribo-depletion libraries. Among the repeat reads, poly(A) samples have the highest fraction of reads mapped to Alu elements, and ribo-depleted samples have the highest fraction mapped to L1 elements (Supplemental Fig. S5). Among the GTEx tissues, testis showed significantly higher expression of SVA F retrotransposons compared to other GTEx tissues (p = 2.46×10^−33^) (Supplemental Figure S6). Furthermore, we observe high co-expression of *Alu* elements and L1 elements across GTEx tissues (R^2^ = 0.7615) (Supplemental Figure 7).

### ROP identifies hyper edited reads

Using standard read mapping approaches, some human reads may remain unmapped due to “hyper editing.” An extremely common post-transcriptional modification of RNA transcripts in human is A-to-I RNA editing (Bazak et al., 2014). Adenosine deaminases acting on RNA (ADARs) proteins can modify a genetically encoded adenosine (A) into an inosine (I). Inosine is read by the cellular machinery as a guanosine (G), and, in turn, sequencing of inosine results in G where the corresponding DNA sequencing reads A. Current methods to detect A-to-I editing sites are based on the alignment of RNA-Seq reads to the genome to identify such A-to-G mismatches. Reads with excessive (‘hyper’) editing are usually rejected by standard alignment methods. In this case, many A-to-G mismatches obscure their genomic origin.

We have identified hyper-edited reads by using the pipeline proposed in (Porath, Carmi, & Levanon, 2014). This hyper-editing pipeline transforms all As into Gs, in both the unmapped reads and the reference genome, and the pipeline realigns the transformed RNA-Seq reads and the transformed reference genome. The method then recovers original sequences and searches for dense clusters of A-to-G mismatches.

A total of 201,676,069 hyper-edited reads were identified across all samples from the three studies. As a control for the detection, we calculated the prevalence of all 6 possible nucleotide substitutions and found that 79.9% (201,676,069/252,376,867) of the detected reads were A-to-G mismatches (Supplemental Fig. S8). In comparison, the in-house RNA-Seq samples have a 96.1% rate of A-to-G mismatches. This massive over-representation of mismatches strongly suggests that these reads resulted from ADAR mediated RNA editing. However, additional experiments are required to confirm the nature of these edits. In addition, we found that the nucleotide sequence context of the detected editing sites complies with the typical sequence motif of ADAR targets (Supplemental Fig. S9).

### The ROP protocol complements transcriptome profiling by non-co-linear RNAs

The ROP protocol is able to detect ‘non-co-linear’ reads via Tophat-Fusion (Kim & Salzberg, 2011) and CIRCexplorer2 (Zhang et al., 2016) tools from three classes of events: (1) reads spliced distantly on the same chromosome supporting trans-splicing events; (2) reads spliced across different chromosomes supporting gene fusion events; and (3) reads spliced in a head-to-tail configuration supporting circRNAs. On average, we observed 816 trans-splicing events, 7510 fusion events, and 930 circular events per individual sample supported by more than one read. Over 90% of non-co-linear events were supported by fewer than 10 samples (Supplemental Fig. S10). We used a liberal threshold, based on number of reads and individuals, because our interest is mapping all reads. However, a more stringent cut off is recommended for confident identification of non-co-linear events, specially in the clinical settings.

Based on the in-house RNA-Seq data, we observe that the library preparation technique strongly affects the capture rate of non-co-linear transcripts. To compare the number of NCL events, we sub-sampled unmapped reads to 4,985,914 for each sample, which corresponded to the sample with the smallest number of unmapped reads among in-house RNA-Seq samples. We observed an average increase of 92% of circRNAs in samples prepared by ribo-depletion compared to poly(A) protocol (p-value = 3 × 10^−12^) (Supplemental Fig. S11). At the same time, we observed an average 43% decrease of trans-splicing and fusion events in samples prepared by ribo-depletion compared to poly(A) protocol (p-value < 8 × 10^−4^) (Supplemental Fig. S11). However, because the tissues differed between protocols (e.g., nasal versus large airway epithelium), this effect might be due in part to tissue differences in NCL events. We view the tissue differences effect to be unlikely. We previously showed that gene expression profiles of nasal airway tissue largely recapitulate expression profiles in the large airway epithelium tissue (Poole et al., 2014).

Furthermore, many NCL events will not be captured by poly-A selection. Therefore, we expect systematic differences in NCL abundance between capture methods. There were no statistically significant differences (p-value > 5×10^−3^) between NCL events in cases and controls. We have compared number of NCL reads across GTEx tissue, and we observe the highest fraction of NCL reads across pancreas samples with 0.75% of reads classified as NCL reads. In all other tissue types, ROP classified approximately .3% reads as NCL reads (Supplemental Figure S12).

### ROP identifies microbial and immune reads and differentiate tissue types and disease status

Reads mapped to B and T cell receptor loci and unmapped reads were used to survey the human adaptive immune repertoires in health and disease. We first used the mapped reads to extract reads entirely aligned to BCR and TCR genes. Using IgBlast (Ye et al., 2013), we identified unmapped reads with extensive somatic hyper mutations (SHM) and reads arising from V(D)J recombination. After we identified all the reads with the human origin, we detected microbial reads by mapping the remaining reads onto microbial reference genomes and phylogenetic marker genes. Here, the total number of microbial reads obtained from the sample is used to estimate microbial load. We use MetaPhlAn2 (Truong et al., 2015) to assign reads on microbial marker genes and determine the taxonomic composition of the microbial communities.

Using in-house RNA-Seq data, we compare immunological and microbial profiles across asthmatics and unaffected controls for the peripheral blood, nasal, and large airway epithelium tissues. A total of 339 bacterial taxa were assigned with Metaphlan2 (Truong et al., 2015) across all studies and are freely available at https://smangul1.github.io/recycle-RNA-seq/.

Using Metaphlan2, we detected bacterial reads in all GTEx tissues except testis, adrenal gland, heart, brain, and nerve. We also observe no bacteria reads in the following cell lines: EBV-transformed lymphocytes(LCLs), Cells-Leukemia (CML), and Cells-Transformed fibroblasts cell lines. On average, we observe 1.43 +−0.43 phyla assigned per sample. All samples were dominated by Proteobacteria (relative genomic abundance of 73%+-28%). Other phyla detected included Acidobacteria, Actinobacteria, Bacteroidetes, Cyanobacteria, Fusobacteria, and Firmicutes. Consistent with previous studies, we observe the nasal epithelium is dominated by Actinobacteria phyla (particularly the *Propionibacterium* genus) (Yan et al., 2013), and the large airway epithelium is dominated by Proteobacteria phyla (Beck, Young, & Huffnagle, 2012) (Supplemental Table S3). As a positive control for virus detection, we used GTEx samples from EBV-transformed lymphoblastoid cell lines (LCLs). ROP detected EBV virus across all LCLs samples. An example of a coverage profile of EBV virus for one of the LCLs samples is presented in Supplemental Fig. S13.

We assess combinatorial diversity of the B and T cell receptor repertoires by examining the recombination of the of Variable (V) and Joining (J) gene segments from the variable region of BCR and TCR loci. We used per sample alpha diversity (Shannon entropy) to incorporate the total number of VJ combinations and their relative proportions into a single diversity metric. We observed a mean alpha diversity of .7 among all the samples for immunoglobulin kappa chain (IGK). Spleen, minor salivary gland, and small intestine (terminal ileum) were the most immune diverse tissue, with corresponding IGK alpha diversity of 86.9, 52.05, and 43.96, respectively (Supplemental Fig. S14-S15). Across all the tissues and samples, we obtained a total of 312 VJ recombinations for IGK chains and 194 VJ recombinations for IGL chains. Inferred recombinations are freely available at https://smangul1.github.io/recycle-RNA-seq/.

Using in-house data, we investigated the effect of different library preparation techniques over the ability to detect B and T cell receptor transcripts. We compared the alpha diversity in large airway samples to nasal samples (Supplemental Fig. S16). Decreased alpha diversity in large airway samples compared to nasal (2.5 for nasal versus 1.0 for large airway) could correspond to an overall decrease in percentage of immune reads. This effect can be attributed to the ribo-depletion protocol not enriching for polyadenylated antibody transcripts. Alternatively, it may result from clonal expansion of certain clonotypes responding to the cognate antigen.

Our comprehensive ROP protocol presents several advantages over previous methods designed to examine features of unmapped reads. First, our method interrogates relationships between features. To explore interactions between the immune system and microbiome, we compared immune diversity against microbial load. Microbes trigger immune responses, eliciting proliferation of antigen-specific lymphocytes. This dramatic expansion skews the antigen receptor repertoire in favor of a few dominant clonotypes and decreases immune diversity (Spreafico et al., 2016). Therefore, we reasoned that antigen receptor diversity in the presence of microbial insults should shrink. In line with our expectation, we observed that combinatorial immune diversity of IGK locus was negatively correlated with the viral load (Pearson coefficient r = −0.55, p-value = 2.4 × 10^−6^), consistent also for bacteria and eukaryotic pathogens across BCR and TCR loci (Supplemental Fig. S17).

Using in-house data, we compared alpha diversity of asthmatic individuals (n = 9) and healthy controls (n =10). The combinatorial profiles of B and T cell receptors in blood and large airway tissue provide no differentiation between case control statuses. Among nasal samples, we observed decreased alpha diversity for asthmatic individuals relative to healthy controls (p-value = 10^−3^) (Fig. 2.b). Additionally, we used beta diversity (Sørensen–Dice index) to measure compositional similarities between samples, including gain or loss of VJ combinations of IGK locus. We observed higher beta diversity corresponding to a lower level of similarity across the nasal samples of asthmatic individuals in comparison to samples from unaffected controls (Fig. 2.c, p-value < 3.7×10^− 13^). Moreover, nasal samples of unaffected controls are significantly more similar than samples from the asthmatic individuals (Fig. 2.c, p-value < 2.5×10^−9^). Recombination profiles of immunoglobulin lambda locus (IGL) and T cell receptor beta and gamma (TCRB and TCRG) loci yielded a similar pattern of decreased beta diversity across nasal samples of asthmatic individuals (Supplemental Fig. S18-S20). Together the results demonstrate the ability of ROP to interrogate additional features of the immune system without the expense of additional TCR/BCR sequencing.

**Figure 3.**
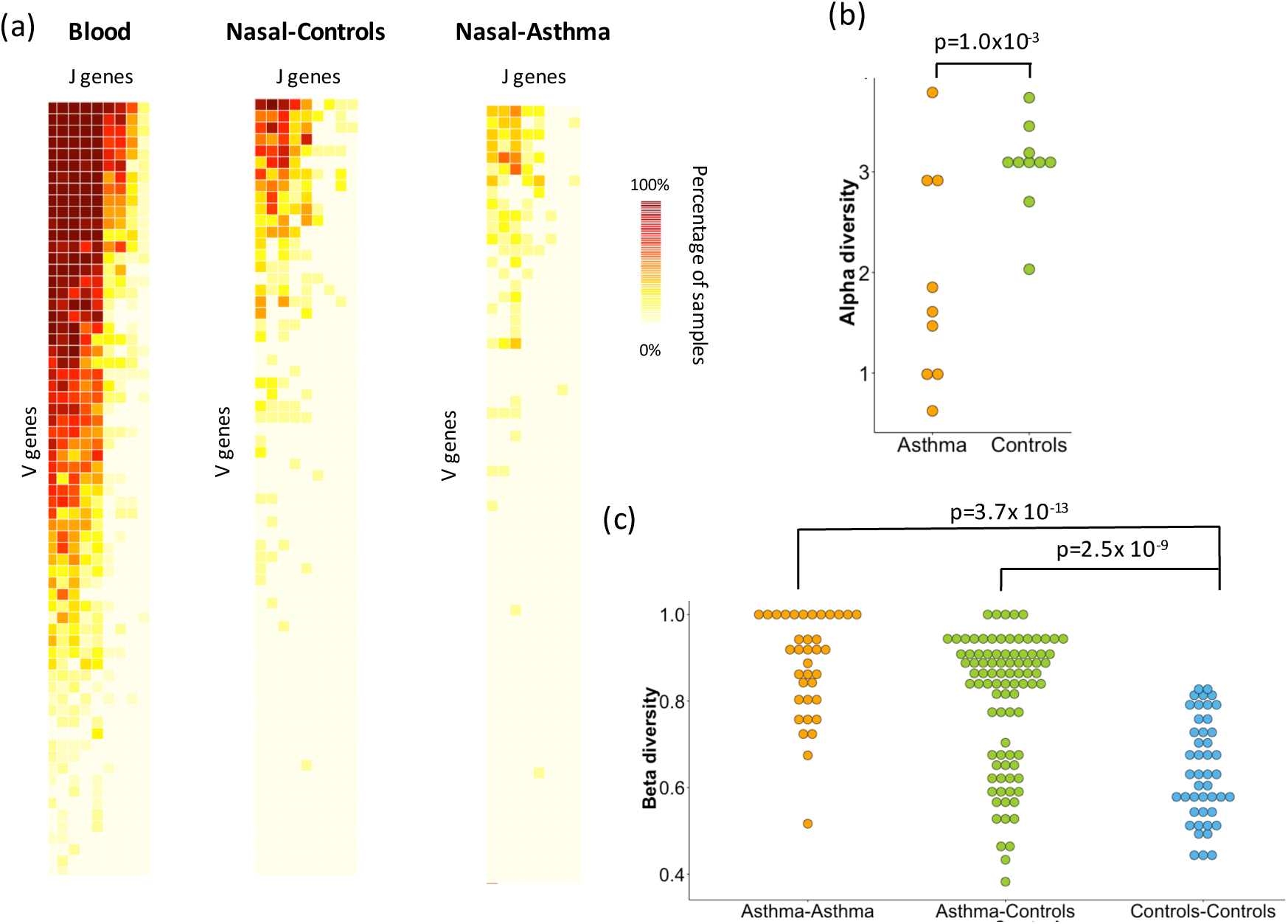
Combinatorial diversity of immunoglobulin kappa locus (IGK) locus differentiates disease status. (a) Heatmap depicting the percentage of RNA-Seq samples supporting of particular VJ combination for whole blood (n = 19), nasal epithelium of healthy controls (n = 10), and asthmatic individuals (n = 9). Each row corresponds to a V gene, and each column correspond to a J gene. (b) Alpha diversity of nasal samples is measured using the Shannon entropy and incorporates total number of VJ combinations and their relative proportions. Nasal epithelium of asthmatic individuals exhibits decreased combinatorial diversity of IGK locus compared to healthy controls (p-value = 1 × 10^−3^). (c) Compositional similarities between the nasal samples in terms of gain or loss of VJ combinations of IGK locus are measured across paired samples from the same group (Asthma, Controls) and paired samples from different groups (Asthma versus Controls) using Sørensen–Dice index. Lower level of similarity is observed between nasal samples of asthmatic individuals compared to unaffected controls (p-value < 7.3 × 10^−13^). Nasal samples of unaffected controls are more similar to each other than to the asthmatic individuals (p-value < 2.5 × 10^−9^).

## DISCUSSION

Our study is the first that systematically accounts for almost all reads, totaling one trillion, available via three RNA-seq datasets. We demonstrate the value of analyzing unmapped reads present in the RNA-seq data to study the non-co-linear, RNA editing, immunological, and microbiome profiles of a tissue. We developed a new tool (ROP) that accounts for 99.9% of the reads, a substantial increase compared over the 82.2% of reads account for using conventional protocols. We found that the majority of *unmapped reads* are human in origin and from diverse sources, including repetitive elements, A-to-I RNA editing, circular RNAs, gene fusions, trans-splicing, and recombined B and T cell receptor sequences. In addition to those derived from human RNA, many reads were microbial in origin and often occurred in numbers sufficiently large to study the taxonomic composition of microbial communities in the tissue type represented by the sample.

We found that both unmapped human reads and reads with microbial origins are useful for differentiating between type of tissue and status of disease. For example, we found that the immune profiles of asthmatic individuals have decreased immune diversity when compared to those of controls. Further, we used our method to show that immune diversity is inversely correlated with microbial load. This case study highlights the potential for producing novel discoveries, when the information in RNA-seq data is fully leveraged by incorporating the analysis of unmapped reads, without need for additional TCR/BCR or microbiome sequencing. The ROP profile of unmapped reads output by our method is not limited to RNA-Seq technology and may apply to whole-exome and whole-genome sequencing. We anticipate that ROP profiling will have broad future applications in studies involving different tissue and disease types.

We observed large effects when using different library preparation protocols on non-co-linear, immunological, and microbial profiles. For example, the poly(A) protocol better captures antibody transcripts by enriching for polyadenylated transcripts, while ribo-depletion protocols capture more circRNAs. The results presented here suggest that selection of a protocol impacts quality of analysis results, and our study may guide the choice of protocol depending on the features of interest.

The ROP protocol facilitates a simultaneous study of immune systems and microbial communities, and this novel method advances our understanding of the functional, interrelated mechanisms driving the immune system, microbiome, human gene expression, and disease etiology. We hope that future efforts will provide a quantitative and qualitative assessment of the immune and microbial components of disease across various tissues. Recent increase in read length and sequencing efficiency provides opportunity for studying individual microbial species and full TCR/BCR sequencing.

## METHODS

### In-house RNA-Seq data

For poly(A) selected samples (n=38), we used a subset of Puerto Rican Islanders recruited as part of the on-going Genes-environments & Admixture in Latino Americans study (GALA II) (Anders, Pyl, & Huber, 2014; Jin, Tam, Paniagua, & Hammell, 2015; Melé et al., 2015; Tarailo-Graovac & Chen, 2009). Nasal epithelial cells were collected from behind the inferior turbinate with a cytology brush using a nasal illuminator. Whole blood was collected using PAXgene RNA blood tubes. RNA was isolated using PAXgene RNA blood extraction kits. For ribo-depleted samples (n=49), we recruited adults aged 18-70 years to undergo research bronchoscopy. During bronchoscopy airway epithelial brushings, samples were obtained from 3^rd^-4^th^ generation bronchi. RNA was extracted from the epithelial brushing samples using the Qiagen RNeasy mini-kit.

Poly(A) selected RNA-seq libraries (n=38) were constructed using 500 ng of blood and nasal airway epithelial total RNA from 9 atopic asthmatics and 10 non-atopic controls. Libraries were constructed and barcoded with the Illumina TruSeq RNA Sample Preparation v2 protocol. Barcoded nasal airway RNA-seq libraries from each of the 19 subjects were pooled and sequenced as 2 × 100bp paired-end reads across two flow cells of an Illumina HiSeq 2000. Barcoded blood RNA-seq libraries from each of the 19 subjects were pooled and sequenced as 2 × 100bp paired end reads across 4 lanes of an Illumina Hiseq 2000 flow cell. Ribo-depleted RNA-seq libraries (n=38), were constructed using 100ng of isolated RNA of large airway epithelium total RNA from 61 samples. Libraries were constructed and barcoded with the TruSeq Stranded Total RNA using a Ribo-Zero Human/Mouse/Rat library preparation kit, per manufacturer’s protocol. Barcoded bronchial epithelial RNA-seq libraries were multiplexed and sequenced as 2 x 100bp paired end reads on an Illumina HiSeq 2500. We excluded 12 samples from further analyses due to high ribosomal RNA read counts (library preparation failure), leaving a total of 49 samples suitable for further analyses.

### GTEx RNA-Seq data

We used RNA-Sequencing data from Genotype-Tissue Expression study (GTEx Consortium v.6) corresponding to 8,555 samples collected from 544 individuals from 53 tissues obtained from Genotype-Tissue Expression study (GTEx v6). RNA-Seq data is from Illumina HiSeq sequencing of 75 bp paired-end reads. The data was derived from 38 solid organ tissues, 11 brain subregions, whole blood, and three cell lines of postmortem donors. The collected samples are from adults matched for age across males and females. We downloaded the mapped and unmapped reads in BAM format from dbGap (http://www.ncbi.nlm.nih.gov/gap).

### SRA RNA-Seq data

Samples (n=2000) were randomly selected using SQLite database from R/Bioconductor package SRAdb (https://bioconductor.org/packages/release/bioc/html/SRAdb.html). We have used a script from https://github.com/nellore/runs/blob/master/sra/define_and_get_fields_SRA.R to select run_accessions from the sra table with platform = ’ILLUMINA’, library_strategy = ’RNA-Seq’, and taxon_id = 9606 (human).

### Workflow to categorize mapped reads

We mapped reads onto the human transcriptome (Ensembl GRCh37) and genome reference (Ensembl hg19) using TopHat2 (v 2.0.13) with the default parameters. Tophat2 was supplied with a set of known transcripts (as a GTF formatted file, Ensembl GRCh37) using –G option. The mapped reads of each sample are stored in a binary format (.bam). ROP (gprofile.py) categorizes the reads into genomic categories (junction read, CDS, intron, UTR3, UTR5, introns, inter-genic read, deep a deep inter-genic read, mitochondrial read, and multi-mapped read) based on their compatibility with the features defined by Ensembl (GRCh37) gene annotations. ROP (rprofile.py) categorizes reads into repeat elements (classes and families) based on their compatibility with repeat instances defined by RepeatMasker annotation (Repeatmasker v3.3, Repeat Library 20120124). We count the number of reads overlapping variable(V), diversity (D), joining (J), and constant (C) gene segments of B cell receptor (BCR) and T cell receptor (TCR) loci using htseq-count (HTSeq v0.6.1).

### Workflow to categorize unmapped reads

We first converted the unmapped reads saved by TopHat2 from a BAM file into a FASTQ file (using samtools with bam2fq option). The FASTQ file of unmapped reads contains full read pairs (both ends of a read pair were unmapped) and discordant read pairs (one read end was mapped while the other end was unmapped). We disregarded the pairing information of the unmapped reads and categorized unmapped reads using the protocol’s seven steps. Reads identified at each step are filtered out.

#### A. Quality Control

Low quality reads, defined as reads that have quality lower than 30 in at least 75% of their base pairs, were identified by FASTX (v 0.0.13). Low complexity reads, defined as reads with sequences of consecutive repetitive nucleotides, were identified by SEQCLEAN. As a part of the quality control, we also excluded unmapped reads mapped onto the rRNA repeat sequence (HSU13369 Human ribosomal DNA complete repeating unit) (BLAST+ 2.2.30).

#### B. Remap to human references

We remapped the remaining unmapped reads to the human reference genome (hg19) and transcriptome (known transcripts, Ensembl GRCh37) using Megablast (BLAST+ 2.2.30). ROP step 3.

#### C. Hyper-editing detection

We used a hyper-editing pipeline (HE-pipeline http://levanonlab.ls.biu.ac.il/resources/zip), which is capable of identifying hyper-edited reads.

#### D. Map to repeat sequences

The remaining unmapped reads were mapped to the reference repeat sequences using Megablast (BLAST+ 2.2.30). The reference repeat sequences were downloaded from Repbase v20.07 (http://www.girinst.org/repbase/). Human repeat elements (humrep.ref and humsub.ref) were merged into a single reference.

#### E. Non-co-linear (NCL) RNA profiling

NCL events include three classes of events: reads supporting trans-splicing events that are spliced distantly on the same chromosome; reads supporting gene fusion events that are spliced across different chromosomes; and reads supporting circRNAs that are spliced in a head-to-tail configuration. To distinguish between these three categories, we use circExplorer2 (v2.0.13). CircExplorer2 relies on Tophat-Fusion (v2.0.13, bowtie1 v0.12.) and allows simultaneous monitoring of NCL events in the same run. To extract trans-spicing and gene fusion events from the TopHat-Fusion output, we ran a ruby custom script, which is part of the ROP pipeline (NCL.rb).

#### D. B and T lymphocytes profiling

We used IgBLAST (v. 1.4.0) with a stringent e-value threshold (e-value < 10^-20^) to map the remaining unmapped reads onto the V(D)J gene segments of the of the B cell receptor (BCR) and T cell receptor (TCR) loci. Gene segments of B cell receptors (BCR) and T cell receptors (TCR) were imported from IMGT (International ImMunoGeneTics information system). IMGT database contains: variable (V) gene segments; diversity (D) gene segments; and joining (J) gene segments.

#### E. Microbiome profiling

We used Megablast (BLAST+ 2.2.30) to align remaining unmapped reads onto the collection of bacterial, viral, and eukaryotic reference genomes. Bacterial and viral genomes were downloaded from NCBI (ftp://ftp.ncbi.nih.gov/). Genomes of eukaryotic pathogens were downloaded from EuPathDB (http://eupathdb.org/eupathdb/). We used MetaPhlAn2 (Metagenomic Phylogenetic Analysis, v 2.0) to obtain the taxonomic profile of microbial communities present in the sample.

### Reference databases

A detailed description of reference databases used by ROP is provided in Supplemental Materials.

### Comparing diversity across groups

First, we sub-sampled unmapped reads by only including reads corresponding to a sample with the smallest number of unmapped reads. Diversity within a sample was assessed using the richness and alpha diversity indices. Richness was defined as a total number of distinct events in a sample. We used Shannon Index (SI), incorporating richness and evenness components, to compute alpha diversity, which is calculated as follows:

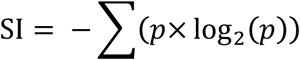

We used beta diversity (Sørensen–Dice index) to measure compositional similarities between the samples in terms of gain or loss in events. We calculated the beta diversity for each combination of the samples, and we produced a matrix of all pairwise sample dissimilarities. The Sørensen–Dice beta diversity index is measured as 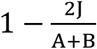, where J is the number of shared events, while A and B are the total number of events for each sample, respectively.

### The robustness of the ROP results against changing the thresholds for each of the ROP steps

We have performed the robustness analysis to investigate the impact of the thresholds used in each step of the ROP approach. For each ROP step, we reported the number of reads identified under different thresholds. The results are presented as cumulative frequency plots (Supplemental Methods Figure SM1).

### The impact of mapping parameters and RNA-Seq aligners on the number of unmapped reads

We investigated the impact of alternative aligners (STAR, https://github.com/alexdobin/STAR) and carefully adjusted the mapping setting to achieve sensitive and very sensitive settings (Supplemental Table S4). The average runtime on Hoffman2 Cluster for Tophat per million reads was 2.5 hours; STAR, 0.13 hours; and Novoalign, 9.1 hours. Novoalign was not considered in the analysis due to its substantially longer running time, which made it infeasible for the protocol.

### Software availability

The ROP software is publicly available at https://github.com/smangul1/rop. The source code for ROP v1.0.4 is also available as Supplemental Material. Custom scripts necessary to reproduce the results and reference files are distributed with Supplemental Material, and are also available with ROP software. A tutorial with detailed instructions on how to run ROP is freely available at https://github.com/smangul1/rop/wiki. For a quick start, an example with 2508 unmapped reads is distributed with the ROP package. Reads are randomly selected from a publically available normal skin (SRR1146076) RNA-Seq sample and might not represent the typical reads of RNA-Seq experiment. The reads are provided for demonstration purposes and are distributed with ROP software. Additional details of ROP, including all parameters and thresholds used, are provided in the Supplemental Methods.

## ACKNOWLEDGEMENT

### DISCLOSURE

The authors declare no competing interests.

### DATA ACCESS

Portions of the nasal airway epithelial whole transcriptome data were published in a previous manuscript (Tarailo-Graovac & Chen, 2009)

